# GraphPred: An approach to predict multiple DNA motifs from ATAC-seq data using graph neural network and coexisting probability

**DOI:** 10.1101/2022.05.02.490240

**Authors:** Shuangquan Zhang, Lili Yang, Xiaotian Wu, Nan Sheng, Yuan Fu, Anjun Ma, Yan Wang

## Abstract

Assay for Transposase-Accessible Chromatin sequencing (ATAC-seq) utilizes hyperactive Tn5 transposase to cut open chromatin and reveal chromatin accessibility at a genome-wide level. ATAC-seq can reveal more kinds of transcription factor binding regions than Chromatin immunoprecipitation sequencing (ChIP-seq) and DNase I hypersensitive sites sequencing (DNase-seq). Transcription factor binding sites (TFBSs) prediction is a crucial step to reveal the functions of TFs from the high throughput sequencing data. TFBSs of the same TF tend to be conserved in the sequence level, which is named motif. Several deep learning models based on the convolutional neural networks are used to find motifs from ATAC-seq data. However, these methods didn’t take into account that multiple TFs bind to a given sequence and the probability that a fragment of a given sequence is a TFBS. To find binding sites of multiple TFs, we developed a novel GNN model named GraphPred for TFBSs prediction and finding multiple motifs using the coexisting probability of k-mers. In the light of the experiment results, GraphPred can find more and higher quality motifs from 88 ATAC-seq datasets than comparison tools. Meanwhile, GraphPred achieved an area of eight metrics radar (AEMR) score of 2.31.

## 1 Introduction

Assay for Transposase-Accessible Chromatin sequencing (ATAC-seq) can recognize and cleave DNA sequences in open chromatin regions via the DNase-I and Tn5 [1]. ATAC-seq obtains more kinds of transcription factor binding regions than Chromatin immunoprecipitation sequencing (ChIP-seq) and DNase I hypersensitive sites sequencing (DNase-seq). The sites of Transcription factors (TFs) binding to DNA sequences are called transcription factor binding sites (TFBSs), which are usually short DNA sequences [2]. The aligned TFBSs of the same TF is the regulatory motif, which can be represented by the position weight matrix (PWM) [3, 4]. So there may be multiple motifs in ATAC-seq data. TFs act as crucial roles in gene expression and human diseases via binding to the DNA sequences [5, 6]. The transcription regulation is carried out by the interplay between TFs and TFBSs, identifying TFBSs aids us to reveal the functions of TFs and the reason for the diseases varied TFs lead to [7]. The footprinting is a fragment of DNA sequence that has not been cleaved due to the transcription factors binding, which can reveal the TFBSs in the genome [8, 9]. Several tools have been developed to detect footprinting from ATAC-seq data, which are based on multi-omics data and hidden Markov algorithm [1, 10]. ATAC-seq footprinting has been applied to TF network prediction [11], comparison of TF activity [10], and identifying TFs enriched in PBMC-specific peaks [12].

In recent years, convolutional neural networks (CNNs) [13] and recurrent neural networks (RNNs) [14] have achieved success in bioinformatics, such as protein-protein networks [15], gene regulatory networks [16], and motif finding [17-19]. The existing DL tools such as scFAN, FactorNet, and DeepATAC employed CNNs to predict TFBSs and find motifs from ATAC-seq data [20-22]. These DL tools utilize convolutional kernels as motif detectors and each motif detector scan input sequences. Concerning ATAC-seq data, a sequence can be bound by multiple TFs, the existing DL model only finds a TFBS from a given sequence by motif detectors.

In a related line of research, graph neural networks (GNNs) [23] can learn the key massage from the graph-structured data. GNNs are seen as the extension of CNNs, which have been shown to obtain excellent performances on molecular structures [24], protein-protein networks [25], gene-gene networks [26], binding preferences of RBPs for RNA structures [27], etc. GNNs can learn the representation of nodes via their neighboring nodes and keep the connection of the graph unchanged, and nodes’ representation also can enable graph-based explanation and reasoning. Due to the successful application of GNNs in bioinformatics, GNNs can enable us to find multiple motifs from ATAC-seq.

To address the above problems, we develop a novel GNN model named GraphPred for TFBSs prediction and finding multiple motifs from ATAC-seq data. GraphPred first builds a heterogeneous graph using ATAC-seq data, which includes three kinds of edges (inclusive edges, similarity edges, and coexisting edges) and two kinds of nodes (sequence nodes and k-mer nodes). GraphPred employs the two layers of GNN, which is used to predict whether the sequences are bound by TFs. Finally, GraphPred calculates the coexisting probability of k-mers using the coexisting edges of the heterogeneous graph and finds multiple motifs from an ATAC-seq dataset. The results demonstrate that GraphPred can find more and higher quality motifs from the ATAC-seq data compared to the comparison tools in the state-of-the-art, and achieved the area of eight metrics radar (AEMR) score of 2.31.

The main contributions of this study are listed below.

1. We build a heterogeneous graph by dividing the given sequences into k-mers. The heterogeneous graph includes three types of edges and two types of nodes and is decomposed to inclusive graph, similarity graph, and coexisting graph.
2. We develop the GraphPred model for predicting TFBSs from ATAC-seq data. GraphPred model employs two layers of GNNs, the first layer of GraphPred is used to learn the embedding of k-mer nodes, the second layer of GraphPred is used to learn the embedding of sequence nodes.
3. GraphPred finds multiple motifs from ATAC-seq data. GraphPred defines the coexisting probability of k-mer nodes to find multiple TFBSs from given sequences and indicates the found motifs are consistent with experimentally verified motifs.
4. We verify the effectiveness of our method across 88 ATAC-seq datasets, which indicates our method can find more and higher quality motifs than comparison tools.

The rest of the paper is organized as follows. Section 2 introduces related works on motif finding using ATAC-seq. The implementation details of the work are described in Section 3. Experimental results are presented in Section 4. In Section 5, we discuss the pros and cons of our method and future works. Section 6 concludes this work.

## 2. Related works

Motif finding aims to find some conserved TFBSs from the high-throughput sequencing data, such as ChIP-seq, ATAC-seq, and DNase-seq data. However, ATAC-seq can detect the whole open chromatin, which means that it can obtain the multiple binding sites of TFs. Footprints of ATAC-seq data enable the investigation of TFBSs, TOBIAS, and HINT-ATAC tools are developed to detect footprinting of ATAC-seq data. TOBIAS utilizes reads from ATAC-seq, known motifs, and sequence annotation in standard formats as input and corrects the inputting reads by the expected signal track [10]. TOBIAS then calculates a footprint score to measure the possibility that a region is a protein binding. HINT-ATAC is based on hidden Markov models, which uses strand-specific, nucleosome size decomposed, and bias-corrected signals to identify footprints [11].

In early literature, k-mers are used features for modeling the properties of DNA sequences [28]. The gkm-SVM model employs the support vector machine (SVM) and gapped k-mers to predict transcription factor binding [29]. The NoPeak utilizes the k-mer and information content to identify TF-binding motifs from ChIP-seq data [28]. Recently, DL algorithms have had successes in bioinformatics, which have been applied to find motifs [30-32]. Deepbind employs CNN and utilized convolutional kernels to find motifs from ChIP-seq data. Deepbind only contains a convolutional layer, a pooling layer, and a fully connected layer, but it achieves better performance than traditional tools [30]. After that, some DL models also are developed to find motifs from ATAC-seq data. DeepATAC encodes the 200 base pair windows of ATAC-seq peaks and employs the CNNs to learn the features of ATAC-seq peaks [22]. Convolutional kernels of the first layer in trained DeepATAC are utilized to find TFBSs from the DNA sequences, and the TFBSs that DeepATAC finds are aligned as a motif. Another DL model, FactorNet utilizes the same strategy of motif finding as DeepATAC. scFAN also employs CNNs to predict the probability that a TF binds to a given DNA sequence. scFAN utilized bulk ATAC-seq data and ChIP-seq data for training, and actually find motifs from the scATAC data via the convolutional kernels of its first convolutional layer.

## 3. Methods

### 3.1 Data and preprocessing

The 88 human ATAC-seq datasets including 80 cell lines are downloaded from the ENCODE project (Supplementary material Table S1). TOBIAS and HINT-ATAC tools with the default parameters are employed to generate footprints [10, 11]. To reduce the bias, we first utilize TOBIAS to generate footprints and calculate footprints scores of each dataset. All footprints are ranked descend by their scores. Then top-1500 footprints are used to intersect with the footprints generated by HINT-ATAC, and intersected footprints of each dataset are applied in the following analysis.

Each intersected footprint is fixed with 101 bps around its centers by the bedtools [33], which are set as positive samples. Since the TFBSs prediction is a binary classification task, we need to collect the negative samples. To train the GraphPred model, we used shuffled positive sequences as negative samples [34]. We give each of the positive samples a label of ‘1’, and give each negative sample a label of ‘0’. For each dataset, 80% of samples are set as training data, 10% of samples are set as validation data and the resting samples are set as testing data. Then, all sequences in each ATAC-seq experiment are divided into k-mers by the step of 1. The sequences and k-mers will be used to construct the heterogeneous graph.

### 3.2 The GraphPred model

To find multiple motifs from ATAC-seq data, we develop a method named GraphPred using GNN and coexisting probability of k-mers (Fig.1). GraphPred formulates TFBSs prediction as a heterogeneous graph-based semi-supervised learning problem (Fig.1A). GraphPred employs two layers of GNN to learn the embedding of k-mer nodes and the embedding of sequence nodes (Fig.1B) and finds multiple motifs using the coexisting probability of k-mer nodes (Fig.1C).

**Fig.1.**
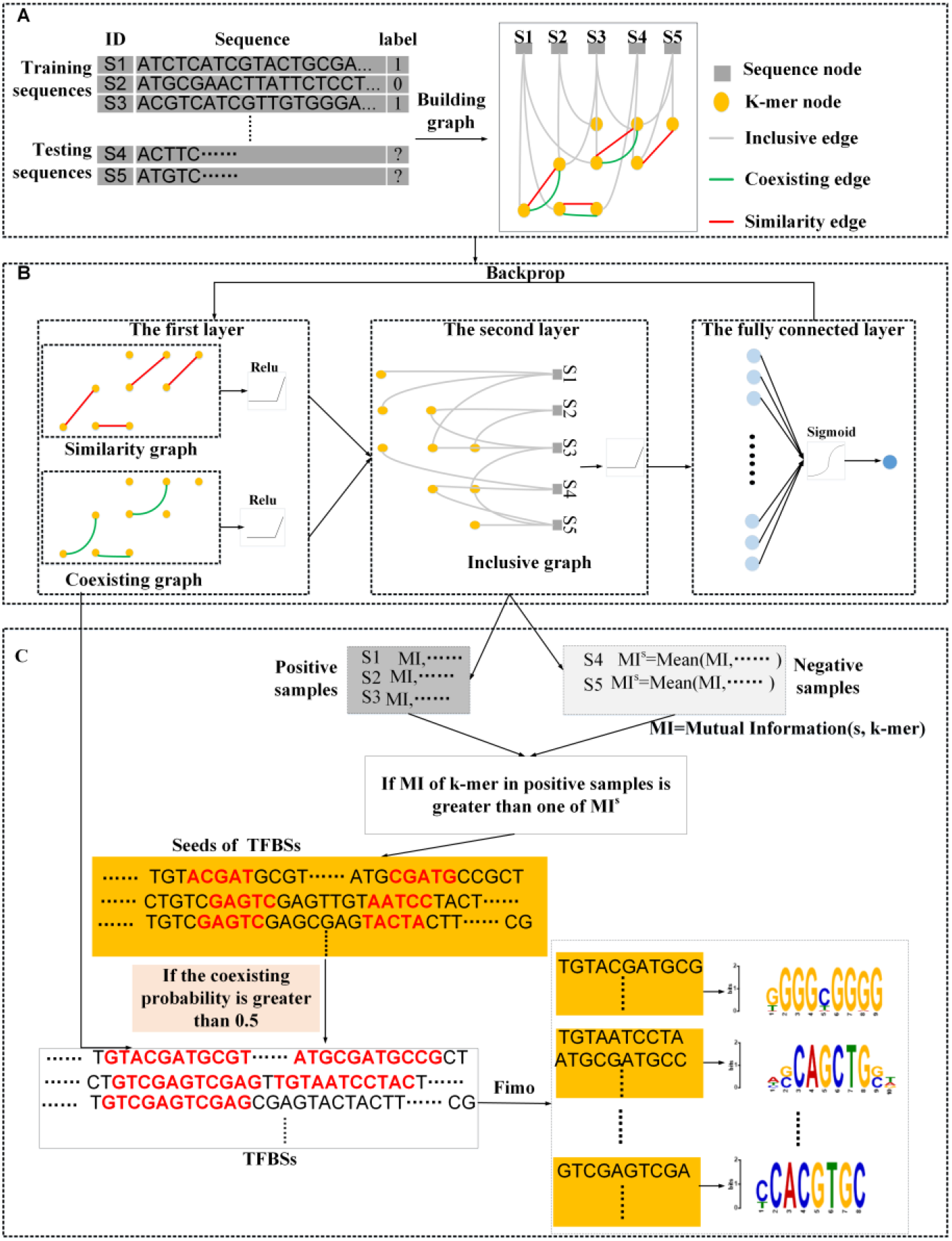
Schematic overview of GraphPred framework. GraphPred builds the heterogeneous graph via given sequences (A). GraphPred learns the representation of sequences to predict the TFBSs (B). GraphPred merges the k-mers to TFBSs using the coexisting probability and finds multiple motifs (C).

### 3.2.1 Building a heterogeneous graph

GraphPred defines three types of edges that connect the k-mer nodes and sequence nodes (Fig.1A). GraphPred first divides the sequences into several fragments of the same length of k by the step of 1, called k-mers. The sequences and k-mers are used to build the heterogeneous graph. Then GraphPred defines inclusive edges, similarity edges, and coexisting edges: 1) an inclusive edge connects a sequence node and a k-mer node if s contains k-mer. Inclusive edges represent the inclusive relations between a given sequence and its constituent k-mers, TF-IDF (term frequency-inverse document frequency) is used to measure the weights of inclusive edges [35]; 2) a similarity edge connects two k-mer nodes of k-mers k1 and k2 by the hamming distance between k1 and k2. Similarity edges represent the relations between two k-mers, the weights of similarity edges are measured by the hamming distances [36]; 3) a coexisting edge connects two k-mer nodes of k-mers k1 and k2 if k1 and k2 tend to coexist in the same sequence. Coexisting edges represent the relations between two k-mers which tend to coexist in sequences.

The three types of edges are shown as follows:

1. Similarity edges. Similarity edges measure the hamming distance of two k-mers. The weight of the edge is given by the formula (1).

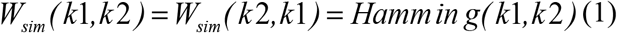

The similarity edges define the mismatch between k-mer nodes in the heterogeneous graph. The two sequence nodes can share their information through k-mers connected by similarity edges, *Hammin g(* ·)m represents hamming distance function, *W*_*sim*_ represents the weight matrix of similarity edges.
2. Coexisting edges. For the given two k-mer *k*1 and *k*2, we define the weight matrix of coexisting edges by *W*_*co*_. We calculate the *W*_*co*_ via the statistical theory, which is given by the formula.

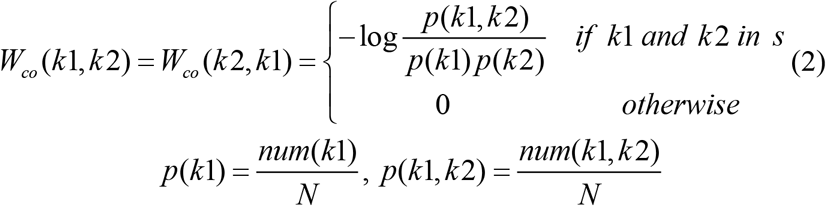

Where *num(* ·) represents the number of the given sequences containing k-mer *k*1 ; *num(k*1,*k* 2 ) is the number of the given sequences containing both *k*1 and *k*2. The log(-) represents the logarithmic function. The definition of coexisting edges enables the information exchange between two k-mers, which coexist in given sequences.
3. Inclusive edges. Each sequence node connects a k-mer node if the given sequence contains the k-mer *k*. We calculate the weights of the edges by the TF-IDF algorithm (formula (3)).

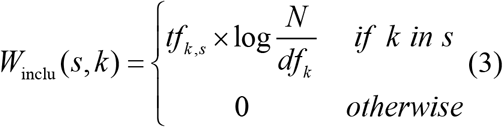

Where *tf*_*k,s*_ is the number of existences of k-mer *k* in sequence *s ; df*_*k*_ is the number of sequences containing *k*, and *N* is the number of sequences including training and testing sequences. *W*_inclu_ represents the weight matrix of inclusive edges.

### 3.2.2 Learning and prediction

We formulate TFBSs prediction as a heterogeneous graph-based semi-supervised learning problem. We decompose the heterogeneous graph into three sub-graphs, i.e coexisting graph, similarity graph, and inclusive graph (Fig.1B). GraphPred employs two layers of GNN to learn the embedding of k-mer nodes and sequence nodes. The first layer of GraphPred is used to learn the embedding of the k-mer nodes from the coexisting graph and similarity graph (Fig.1B). The second layer is used to learn to embed the sequence nodes from the inclusive graph.

We normalize the adjacency matrix based on coexisting graph and similarity graph to obtain k-mer nodes’ original embedding *h*_*co*_ and *h*_*sim*_ respectively.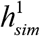 is the simh embedding of k-mer nodes that GraphPred learns from the similarity graph. GraphPred employs GNNs to update each embedding of k-mer nodes by neighbors of k-mer nodes.

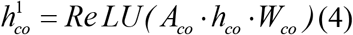

Where ·represents inner product, *Re LU(* ·) represents rectified linear unit function, *W*_*co*_ is the weight of the coexisting layer, *A*_*co*_ is an adjacency matrix based on the coexisting graph, 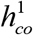 represents the embedding of k-mer nodes based on 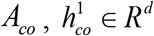, the *d* is the number of dimensions.

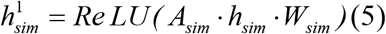

Where*W*_*sim*_ is the weight of the coexisting graph, *A*_*sim*_ is an adjacency matrix based on the coexisting graph, 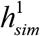 represents the embedding of k-mer nodes based on 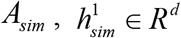, the *d* is the number of dimensions.

Then, the embedding based on 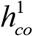 and 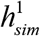 of each k-mer node is merged into a vector *h*_*co _ sim*_ (formula (6)), which is used as the input of the second layer. GraphPred learns the representation of the sequence nodes via the second layer.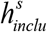 represents the embedding of the sequence nodes in the inclusive graph.

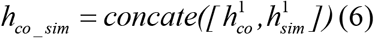

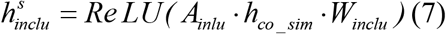

Where 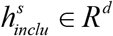

The output layer of GraphPred is a fully connected layer:

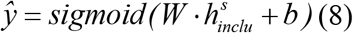

Where*W* represents the weight matrix, *b* represents the bias, *sigmoid(* ·) is a sigmoid function.

We select the Binary Cross Entropy as the loss function (BCELoss) of GraphPred model:

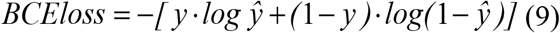

Where *ŷ* represents the output of the GraphPred model, *y* represents the true label.

### 3.2.3 GraphPred finds multiple motifs using the coexisting probability of k-mers

GraphPred predicts whether a given sequence has TFBSs by learning the representation of nodes from the heterogeneous graph. GraphPred calculates the MIs [37] between the embedding of k-mer nodes and sequence nodes (Fig.1C). For each negative sequence, GraphPred calculates *MI*^s^ by averaging MIs between the negative sequence and each k-mer in it (formula (11)). Then GraphPred calculates the MI of each k-mer in a positive sequence (formula (10)). If the MI of a k-mer is greater than one of *MI*^s^, the k-mer comes from a positive sequence is set as a seed of a TFBS. In the above way, GraphPred can find multiple seeds of TFBSs from a given sequence.

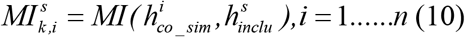

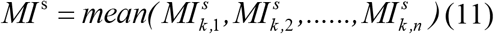

Where *i* is *i* th k-mer, *s* represents a given sequence, *n* is the number of k-mers in a given sequence, the 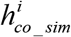 represents the embedding of *i* th k-mer which is given by formula 6, *k* represents the length of a k-mer.

Next, GraphPred finds the location of a seed of a TFBS and defines two k-mers around the center of the position (±5bps). GraphPred calculates a coexisting matrix by the formula 12, each value of the coexisting matrix represents the possibility that two k-mers are bound by a TF (Fig.2). If the coexisting probability of two k-mers is greater than 0.5, two k-mers are merged as a TFBS (formula (13)).

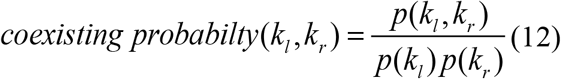

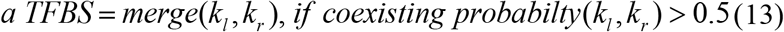

Where *k*_*l*_ represents a k-mer in the left of the center of a TFBS seed, *k*_*r*_ represents a k-mer in the right of the center of a TFBS seed, *coexisting probabilty* represents the coexisting probability of *k*_*l*_ and *k*_*r*_ which are calculated from the coexisting graph.

Finally, GraphPred utilizes Fimo to compute the p-value of each TFBS by querying each motif of the HOCOMOCO V11 database [38]. If the p-value of a TFBS is less than 0.05, it will be seen as a TFBS that a TF binds to.

### 3.3 Evaluation metrics

The precision, recall, F1_score, accuracy (ACC), specificity, Matthews correlation coefficient (MCC), the area under the receiver operating characteristic curve (AUROC), and area under the precision-recall curve (AUPRC) are summed as the area of eight metrics radar (AEMR) score, which is set as an overall score to rank all models. Except for AEMR, other metrics can be computed by the confusion matrix [39], but the AEMR score is a comprehensive metric. Once the eight scores are calculated, a radar chart can be generated that consists of eight equiangular spokes with each spoke representing one of the scores defined above.

We first calculate the precision, recall, F1_score, specificity, ACC, MCC, AUROC, and AUPRC by the confusion matrix, which is shown as below:

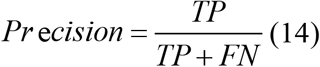

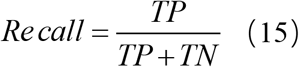

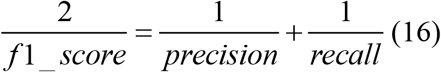

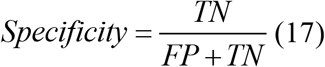

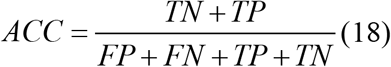

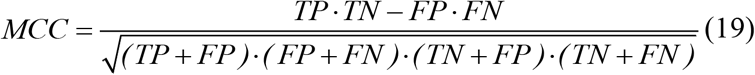

Where TP, TN, FP, FN represent the number of the true positive (TP), true negative (TN), false positive (FP), false negative (FN), respectively.

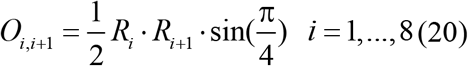

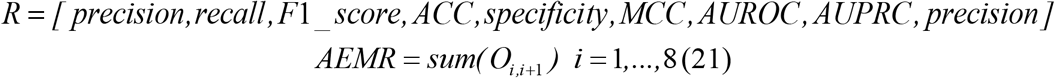

The AEMR score is the total area of the octagonal radar. The higher the AEMR score is, the better performance the tool has for sequence classification.

## 4 Results

### 4.1 Training of the GraphPred model

Before starting to train the GraphPred model, we first represent the k-mer nodes via normalizing the adjacency matrix based on coexisting and similarity edges. After constructing the heterogeneous graph, we use the 150 dimensional vectors to represent the embedding of k-mer nodes and sequence nodes, respectively. We train GraphPred model by the 50 epochs using Adam optimizer [40]. The initial learning rate of GraphPred is 0.001 and is adjusted by the natural exponential decay with 0.001. For each ATAC-seq experiment, we train the GraphPred on the training dataset and test the model on the testing dataset. The dropout of 0.2 is utilized to prevent the overfitting issue. Meanwhile, the length of k-mer is decided by our experiment, the 3-mer, 5-mer, 7-mer, and 9-mer are tried. Considering the computational complexity and the performance of the model, all given sequences are divided into 5-mer.

### 4.2 TFBSs prediction

Our evaluation metrics included precision, recall, F1_score, ACC, specificity, MCC, the AUROC, AUPRC, and AEMR. The AEMRs score of all models on the testing data is shown in Fig. 3. We selected the scFAN, DeepATAC, and FactorNet models as comparison tools in this study. DeepATAC is a DL model to predict TFBSs on the ATAC-seq data, which obtains the AEMR of 1.63 in our experiment (Fig.3B). The scFAN achieved an AMER of 1.51, which is lower than DeepATAC. GraphPred achieved a 2.21 AEMR score, which was better than all comparison models. Our GraphPred model improved the AEMR by 18.2% (Fig.3).

**Fig.3.**
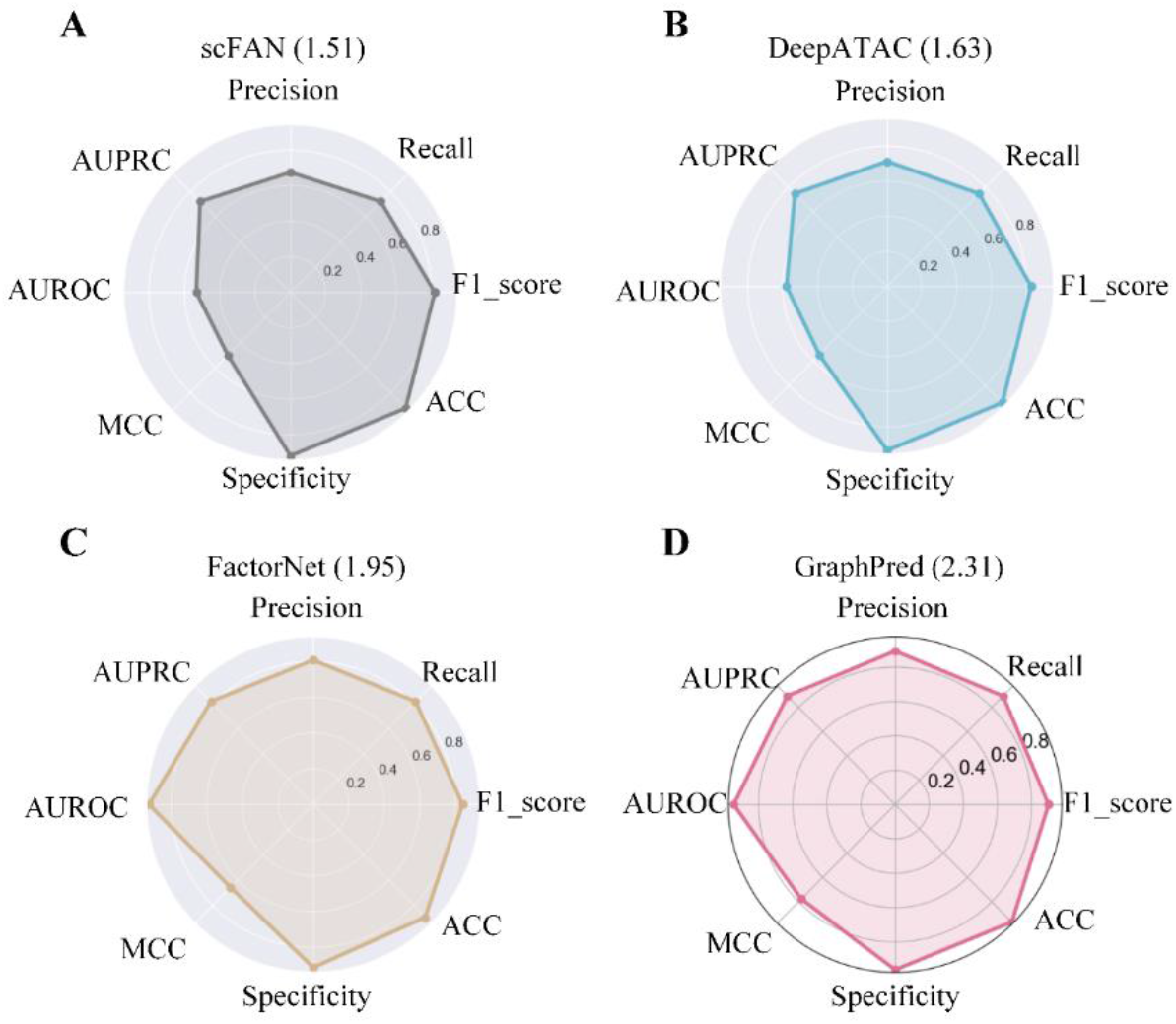
All models obtain average AEMR scores on 88 ATAC-seq datasets. GraphPred model obtained the highest AEMR score.

### 4.3 Finding multiple motifs from ATAC-seq data

To interpret our method, GraphPred calculates MI values between each sequence and each k-mer they contain. The value of MI represents the importance of each k-mer to sequence if k-mers are in a sequence. Then, the coexisting probability of two k-mers is used to find TFBSs, which enables GraphPred to find multiple TFBSs from a given sequence. For each ATAC-seq experiment, we found motifs by the above way and retained motifs that ICs at least three positions are larger than 1. The found motifs are compared to the HOCOMOCO v11 database via the TOMTOM v5.1.0 tool [2, 41]. The matched motifs are significant if their p-values are less than 0.05. Taking the GSE172538 dataset as an example, Fig. 4A shows the ROC and PRC curves of all models, among which GraphPred achieved the highest AUROC and AUPRC scores. As shown in Fig.4B, GraphPred found four significant motifs.

**Fig.4.**
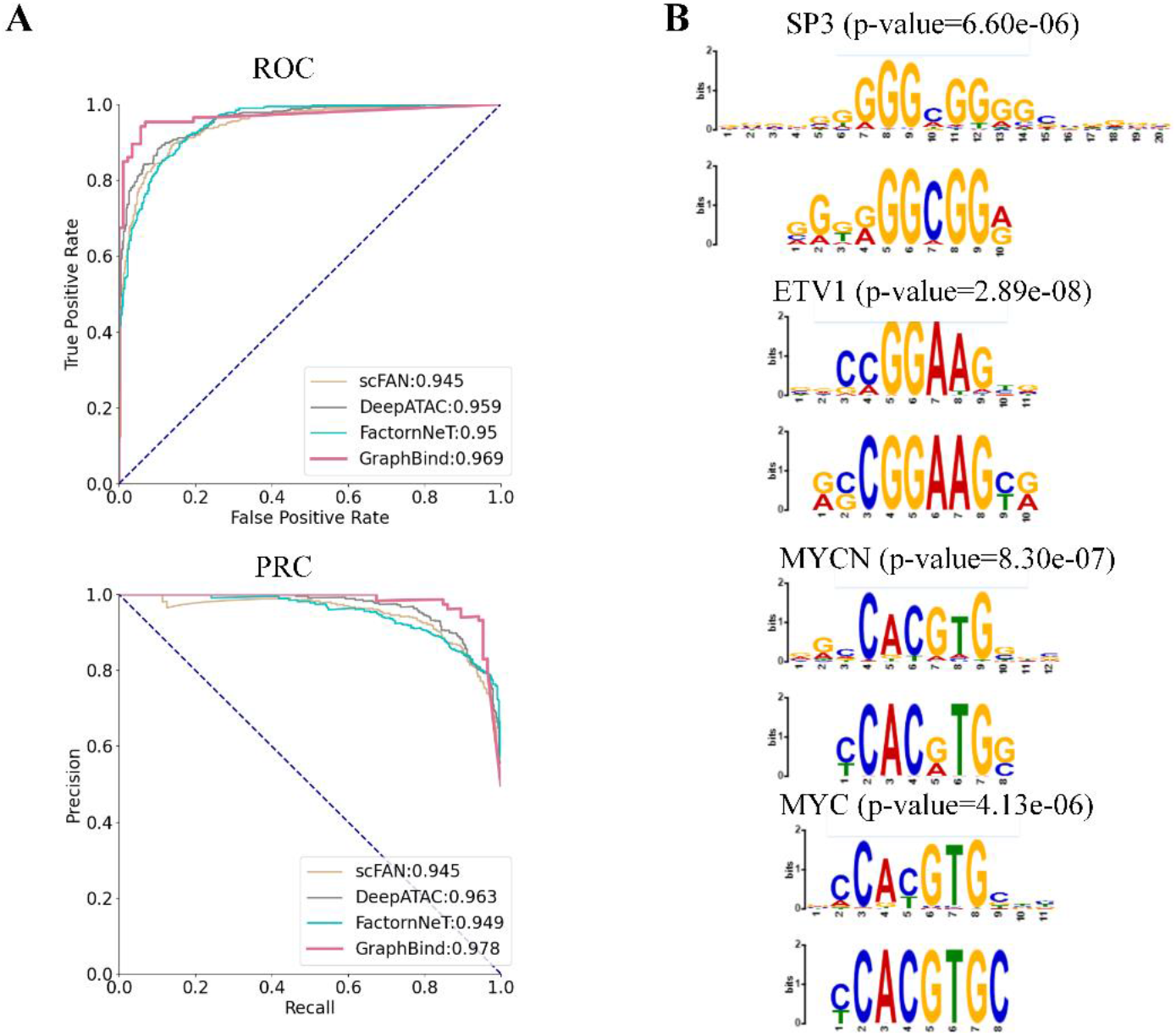
The ROC and PRC curves of all models (left) and the four significant motifs (right) are found by GraphPred. The found motifs (bottom) are matched to the HOCOMOCO V11 database (top) by the TOMTOM.

To validate the ability of GraphPred to find multiple motifs, we use all models to find motifs from 88 ATAC-seq datasets. We employ TOMTOM tool to measure the quality of found motifs by querying the HOCOMOCO database. TOMTOM calculates the p-value to evaluate the quality of found motifs. All found motifs from 88 ATAC-seq datasets were defined as the representative motifs if their p-value is less than 0.05. MEME-chip is a traditional tool to find motifs, which is selected as a comparison tool. Eventually, 337 significant motifs are found from all 88 ATAC-seq datasets for the following analysis. Among five models, our GraphPred model found 207 significant motifs which are more than motifs that comparison models found (Table1). The result of the conventional method MEME-chip was also shown in table 1, MEME-chip only found 55 significant motifs.

**Table1.** Counts of found motifs by each model

| Models | MEME-chip | scFAN | DeepATAC | FactorNet | GraphPred |
| --- | --- | --- | --- | --- | --- |
| Count of motifs | 55 | 199 | 181 | 176 | **207** |

As showcased in Fig.5A, we showed the overlapped and unique status of the identified 337 motifs. Of all 337 found motifs, 10 motifs are only shared by all models. However, only 31 motifs are unique to our GraphPred. To compare the quality of 337 motifs, we show the p-value that each model finds motifs. Showcased in Fig.5B, GraphPred found the highest quality motifs compared to competing tools. Although DeepATAC, FactorNet, and scFAN utilize hundreds of convolutional kernels as motif detectors, they are unable to find higher-quality motifs. Through the above analysis, we showed that our method finds more and higher quality motifs than all comparison tools.

**Fig.5.**
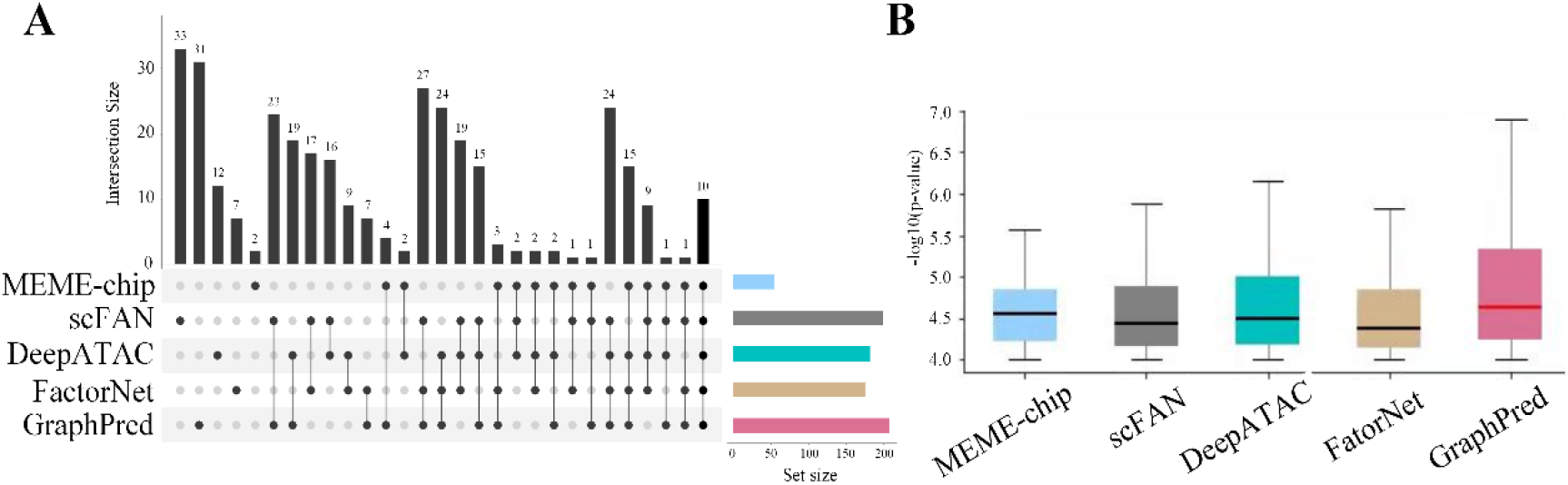
Analysis of motif on 88 ATAC-seq datasets. The shared and unique motifs between the five methods (A). Box plot of motif enrichment P-value of five methods concerning 88 ATAC-seq datasets (B).

## 5 Discussion

In this study, the footprint of ATAC-seq is used to construct the heterogeneous graph. We utilized HINT-ATAC and TOBIAS to find footprints of ATAC-seq which maybe generate false-positive samples. However, our method found more and higher quality motifs, indicating that our method is reliable. GraphPred constructed the heterogeneous graph by dividing the given sequences into k-mer nodes. Nodes of the heterogeneous graph are connected by three types of edges, which show three types of relationships between nodes. However, the positional relationship between k-mer nodes has not been effectively utilized. The position embedding was widely applied in natural language processing[42, 43]. In bioinformatics, the DNA sequence can be seen as a sentence and the base is a word. Our method can utilize position embedding to represent the positional connections between bases of DNA sequences.

We decomposed the heterogeneous graph into the inclusive graph, similarity graph, and coexisting graph. GraphPred employed a two-layer of GNN to predict whether the given sequences are bound by TFs. The first layer was used to learn the embedding of k-mer nodes from the similarity graph and coexisting graph, the second layer of GraphPred was used to learn the embedding of sequence nodes from inclusive graph. Some novel algorithms can be applied in TFBSs prediction, such as graph attention networks [44], graphGAN [45], and graph autoencoders [46]. In the heterogeneous graph, there are many relationships between nodes and each kind of relationship represents the importance between two nodes. The attention mechanism can allocate and update the different weights to the nodes and edges of the heterogeneous graph, in the training process of the model. Our model can capture the important nodes and edges via their weights.

Meanwhile, TFs bind indirectly to motifs of other TFs, which co-regulate targeted gene expression [47]. The cooperation of TFs acts as a vital role in the process of human diseases [48]. ATAC-seq [49] data can detect open accessible DNA regions by probing open chromatin, meaning that ATAC-seq data contains multiple TFs. By analyzing ATAC-seq data, we can reveal the interaction between TFs, and explore the inducement of human disease. Therefore, ATAC-seq provides a potential opportunity to study the cooperation between different TFs.

## 6 Conclusions

In this study, we developed a novel method named GraphPred for finding multiple motifs from ATAC-seq data. GraphPred divided the given sequences into 5-mers for the construction of a heterogeneous graph. The heterogeneous graph contains two types of nodes and three types of edges. We conducted experiments on 88 ATAC-seq datasets to analyze the effectiveness of the proposed method. In particular, the AEMR is an overall score that is used to rank all models. GraphPred found multiple motifs from ATAC-seq data through the coexisting probability of k-mers. Regarding ATAC-seq data, our method found more and higher quality motifs, which demonstrated the method based on coexisting probability of k-mers is more efficient than DL models. As a result, GraphPred achieved better performance than a few state-of-the-art methods. This study makes a great contribution to finding motifs from ATAC-seq data.

## Declarations

### Consent for publication

Not applicable.

### Availability of data and code

The ATAC-seq data can be downloaded via https://www.encodeproject.org/. The code of GraphPred model is freely available at https://github.com/zhangsq06/GraphPred.git

### Authors’ contributions

SZ conceived the project. SZ, XW, NS, and collected the data and performed the experiments. YW and AM designed the study. SZ, LY, and YF wrote the manuscript. All authors read and approved the final manuscript for publication.

