## supplemental files for "GraphPred: An approach to predict multiple DNA motifs from ATAC-seq data using graph neural network and coexisting probability"

### Supplementary materials

Table S1. The GEO number of 88 ATAC-seq datasets

| | | | |
|---|---|---|---|
| GSE114200 | GSE169767 | GSE172977 | GSE172833 |
| GSE114201 | GSE172674 | GSE173067 | GSE172869 |
| GSE114202 | GSE172546 | GSE172538 | GSE172575 |
| GSE114204 | GSE172700 | GSE172638 | GSE172623 |
| GSE114205 | GSE172719 | GSE172752 | GSE172669 |
| GSE169955 | GSE172594 | GSE172760 | GSE172620 |
| GSE170012 | GSE172702 | GSE172814 | GSE172775 |
| GSE170214 | GSE172553 | GSE172559 | GSE172603 |
| GSE170245 | GSE169929 | GSE172679 | GSE172966 |
| GSE170918 | GSE172979 | GSE172646 | GSE172954 |
| GSE170251 | GSE173002 | GSE172772 | GSE172952 |
| GSE170378 | GSE173044 | GSE172949 | GSE172945 |
| GSE172699 | GSE172751 | GSE172990 | GSE172841 |
| GSE170824 | GSE172788 | GSE172986 | GSE172722 |
| GSE170337 | GSE172797 | GSE173023 | GSE172727 |
| GSE172731 | GSE172846 | GSE173027 | GSE172793 |
| GSE169929 | GSE172894 | GSE173043 | GSE172826 |
| GSE172886 | GSE172976 | GSE172904 | GSE172626 |
| GSE172975 | GSE172634 | GSE172934 | GSE172592 |
| GSE172805 | GSE172803 | GSE172959 | GSE172710 |
